# MHC class I / II restricted T cell epitopes from clinical isolate of *Mycobacterium tuberculosis*: A potential candidate for vaccine development for Tuberculosis

**DOI:** 10.1101/2024.09.13.612852

**Authors:** Niharika Sharma, Bhawna Sharma, Beenu Joshi, Santosh Kumar, Keshar Kunja Mohanty, Hridayesh Prakash

## Abstract

Tuberculosis is major challenge to the health care system with TB associated death rates increasing annually. Optimum management of TB (particularly latent or MDR cases) warrants use of immunological approaches like subunit or peptide-based vaccination for tailoring effector immunity in patients. Since MHC class I is a potent enhancer element of host immunity and effective in clearing large variety of intracellular pathogens or tumors. In this context, we explore whether MHC-I restricted peptides from clinical isolates of *M. tuberculosis* can be used as an adjuvant for augmenting host immune responses. In the present study, we have synthesized various peptides from clinical isolates of *M. tuberculosis* which were having high affinity for Class I MHC molecules as potential immune enhancer for T cell or iNKT cell populations. We have evaluated the immunogenic potential of various MHC class I restricted epitopes (Rv2588c, Rv1357, Rv0148, Rv2973, Rv2557 and Rv2445) which were derived from clinical isolates of *M. tuberculosis* on increased proliferation of T or iNKT cells, release of IFN gamma secreted by T cells as well as NO as indicative parameters of immuno-stimulation. As expected, FACS and ELISA data clearly revealed that these peptides were potentially immunogenic for PBMCs from both healthy as well as 10 HC PTB patients. Our data clearly demonstrated a significant immune response in the PBMC from w PTB patients over healthy individuals which mimicked booster response. Our cytokine and nitric oxide data further revealed the influence of these peptides on sensitizing innate immune response as well.

**Significance:** Our study demonstrates the significance of MHC class I restricted peptides from *M. tuberculosis* for inducing potential immunogenic responses in host that may qualify them as potent vaccine candidate. To the best of our knowledge this is the first immune monitoring protocol describing the impact of synthetic novel MHC class I restricted T-cell epitope (Rv2588c, Rv0148) on cell mediated and innate immune response in PBMC populations and suggests their potential as vaccine candidate

## Introduction

TB remains a significant global health burden, with India accounting for a high percentage of both TB cases and TB-related deaths [1]. The study addresses this burden by exploring new strategies for TB immunotherapy. The research focuses on the immunogenicity of MHC class I restricted peptides derived from *M. tuberculosis* proteins, which could potentially enhance the immune response against TB, offering a new approach to vaccine development. The study evaluates the T cell response to these synthetic peptides in pulmonary TB patients and healthy controls, providing insights into the immune mechanisms that could be harnessed for vaccine development. By identifying peptides that elicit a significant immune response, particularly in TB patients, the study suggests that these peptides could serve as potential memory response triggers and new vaccine candidates. The research uses a combination of bioinformatic predictions, ex vivo analysis, and immunological assays to verify the potential of these peptides as immunogenic epitopes, which is a rigorous approach to candidate vaccine evaluation.

Understanding the mechanism of host cell immune response along with the antigenic structure of pathogen is extremely crucial in vaccine development studies. Availability of the entire *M. tuberculosis* genome sequence led to the expedited research for identifying the specific antigens which can help in mounting a protective Th1 response [2]. Various early phase secreted whole-protein antigens of *M. tuberculosis* such as Hsp60, Ag85A, Ag85B, ESAT-6, CFP-10 have already hit the clinical trials for their ability to be recognized by TLRs and stimulate Th1 type cells. But these whole protein-based vaccines provide limited protection as they are only expressed during early stages of active TB infection [3]. Further, they unnecessarily increase the immunogenic load and may lead to allergic responses in the host [4]. Also, neither whole pathogen nor the pathogen whole protein is immunogenic; only specific amino acid fragments within the pathogen acts as potential immunogen and are sufficient to mount protective immune response in the host. This rationale leads to the designing of ‘peptide-based vaccine’ which incorporates only specific immunogenic peptide epitopes. Identification of all the potential immunogenic epitopes within the pathogen genome can be done using Reverse vaccinology approach with the help of available software. Incorporation of antigens expressed during both early and late phase of TB infection with or without BCG could be an ideal approach to design an effective vaccine against *M. tuberculosis*. Thus, entirely synthetic peptide-based vaccines hold a promising future for vaccination [5]. *M. tuberculosis* pathogenesis initially starts with recognition of pathogen associated molecular patterns (PAMPs) by receptors on immune cells especially macrophages and dendritic cells [6]. After recognition, antigens are processed into smaller peptide fragments and mounted on Major Histocompatibility Complex (MHC) molecules. MHC class II loaded with peptide epitope prime the CD4+ helper T cells which further activates the CD8+ cytotoxic T cells. The T cell subsets are known to be crucial in host mediated immune response to pathogen [7]. Macrophage activation by T cells appears to be a central step of acquired resistance against *M. tuberculosis* [8]. T lymphocytes are known to secrete different types of interleukins and pro-inflammatory cytokines like Interferon-γ (IFN-γ), and IL-2 that are involved in the activation of macrophages [9]. IFN-γ activates macrophages and stimulates them to ingest and kill mycobacterium more effectively [10]. The role of CD4+ T cells as chief mediators of anti-tuberculous immunity has been reported through many studies [11].The importance of cytokine mediated activation of macrophages and control of bacterial growth is well studied in mouse models [12] CD8+ T cells also have a cytolytic function which results from recognition of mycobacterium antigens presented by MHC Class I molecules on the surface of infected macrophages [13]. Out of both Class I or II, MHC class I is decisive for antigen presentation by APC / phagocytes to T cells for augmenting immune response against intracellular pathogens and / or tumors. MHC restriction is a critical parameter and paramount for vaccine development and for triggering effector immunity against infectious diseases. Peptide based vaccination often trigger protective immunity against pathogens and for that purpose selection of peptide with high grade homology and / or affinity with MHC plays an important role in triggering protective immune response. Following this hypothesis, we used differentially expressed *M. tuberculosis* peptides from TB patients which were previously purified by mass spectroscopic approach [14] Immunoprofiling method revealed their high immunogenic nature over H37Rv when tested in human PBMC cultures [15]. Bioinformatics and BCPRED tool further screened Rv0148, Rv2588c, Rv0635, Rv0896, and Rv0951 as potent B-cell epitopes in the PBMC from139 PTB cases and 52 healthy individuals. This was later supported by a high titre of antibody against these peptides in PTB patients which further indicated their diagnostic importance for TB [16].

B cell responses are correlative but not sufficient for effective control of TB burden in the host which requires a concerted activation of innate and cell mediated immune compartment. In view of this, the present study investigated the ability of these peptides to augment cell mediated immune response in the PBMC from healthy as well as PTB patients for qualifying as a potential vaccine candidate against TB. For that purpose, we investigated the immunogenicity of Rv0148, Rv2588c, Rv0635, Rv0896, and Rv0951 on T-cell response. Our immune monitoring results potentially suggested that apart from B cells, these peptide enhanced T cell responses in the PBMC from 10 PTB patients and 10 healthy individual. Sub cellular analysis further revealed that these peptides augmented the IFN/ GM-CFS+ CD4/8+ T cells populations in the PBMC of PTB patients. Taken together our study revealed that these peptides are capable of eliciting T cells response as well and they can qualify as a vaccine candidate that elicits a heightened population of IFN-γ/GM-CSF+ CD4/CD8+T cells as compared with the unstimulated PBMC.

## Materials and methods

### Patients’ recruitment

The entire research protocol was approved by the Institute Human Ethics Committee of National JALMA Institute for Leprosy and Other Mycobacterial Diseases (NJIL&OMD), Agra. Written consent was obtained from participants before sample collection. A total of 10 PTB patients with age groups from 18-55 years that were displaying clear clinical symptoms, such as benchmark respiratory problems, positive mucus smear microscopy, culture tests, and radiology of chest, were recruited from the Outpatient Department (OPD) of Sarojini Naidu Medical College Agra as per the scheme shown under **(Figure 1).** Purified protein derivative (PPD) obtained from ARKRAY, India, was inserted into the forearm muscle of individuals as per the guidelines, 0.1 mL of liquid PPD comprised 5TU tuberculin was used. Within 48–72 hours, reading was noted. The presence or absence of the induration was observed. Induration was considered positive if the diameter was above 10 mm. The PPD status of patients was recorded. HIV positive patients with symptoms of other forms of tuberculosis and patients suffering from immunosuppressive diseases like diabetes mellitus were not included in the study. A total of 10 healthy individuals with age group from 18-55 were recruited. Only those healthy who had no history of TB, or any other mycobacterial or infectious diseases were included. The PPD grade of all the participants was considered.

**Figure.1:**
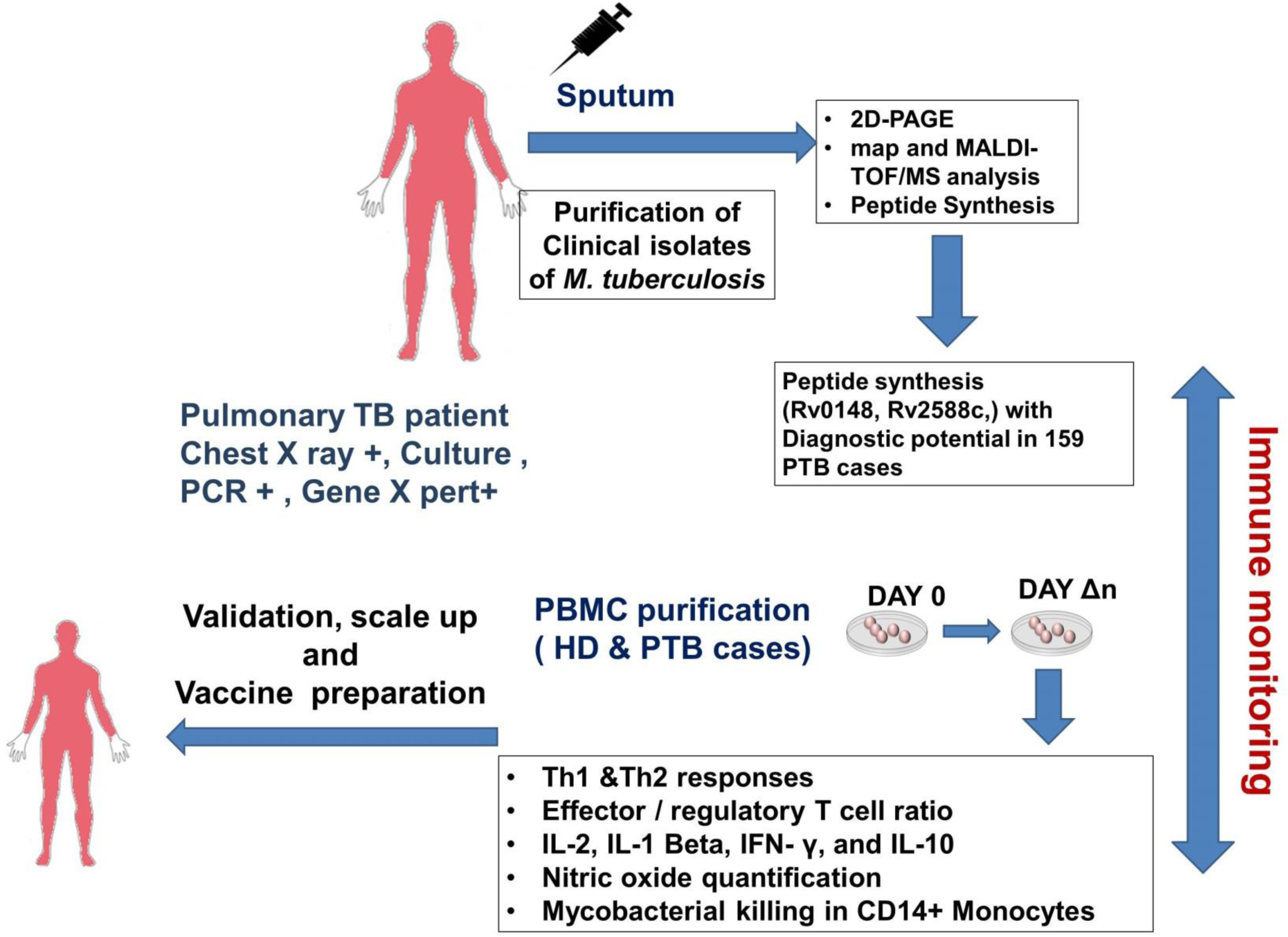
Schematic representation. Scheme adapted for the immuno monitoring of *M.tb* peptide in the PBMC from healthy donors and PTB patients comparatively

### Prediction of T-cell epitope and synthesis of peptides

Peptides Rv1357c (TAFVLEQACRHVRSW), Rv2973 (ALMHGRLSADDKDAA), Rv2588C (IMGGFMYFASRRQRR), Rv2557 (EGWIVYARSTTIQAQ), Rv2445c (LLEFITSGPVVAAIV) and Rv2901c (ADAWVWDMYRPARFV) of *M. tuberculosis* having T-cell epitopes as predicted by bioinformatics tools Propred-2, HLApred, HLA-DR4pred for MHC-II and Propred-1, nHLApred, HLApred for MHC-I were synthesized commercially by GL Beachem, Shanghai Ltd. China according to our specification.

### Immune monitoring of peripheral blood mononuclear cells (PBMCs)

The PBMC from both healthy and TB patients were purified using the ficoll based method. Briefly 2ml ficoll hypaque was taken into a 15ml centrifuge tube. Heparinized blood sample was diluted (1:1) with Rosewell Park memorial institute (RPMI) media. Diluted blood (4ml) was carefully layered on to ficoll hypaque (2ml) in a ratio of 1:2. It was centrifuged at 450*g for 20 minutes at 25^0^C. Lymphocyte layer (white buffy layer) was carefully taken into a fresh 15ml centrifuge tube. At least 3 volume of hank’s balanced salt solution (HBSS) was added to the peripheral blood mononuclear cells (PBMCs) in the centrifuge tube. PBMCs were cultured in U-bottom 96 well plates, stimulated with different doses of peptides having T cell epitopes (2.5, 5, 10μg/ml of PBS) and PPD (5μg/ml) in triplicate and kept for 6 days at 37^0^C in 5% CO_2_ incubator. The viability of these cells was checked by the MTT based method. Supernatant from the PBMC was collected after 24hours, 48 hours and 5 days of culture and IL-1β, IFN-γ, IL-4 and IL-10 were estimated using commercially available kits (R&D System).

### Flow cytometry analysis

FACS staining was done to analyse surface molecules (CD3, CD4, CD25, CD69, CD45Ra, iNKT) and intracellular cytokine screening was done to monitor (IL-2, IFN-γ, IL-4 and IL-10 positive T cells in PBMC cultures. After isolation of PBMCs from an individual, cells were counted, and a cell suspension (10^6^cells/ml) was prepared using 5% cRPMI. Cells were stimulated with peptides along with PPD and incubated at 37^0^C in humidified 5% CO_2_ incubator for 16-18 hours. After incubation 5µl of brefeldin (1:10 diluted in cRPMI) was added in each well and left for incubation for 4-6 hours. After incubation, the plate was centrifuged at 250g for 5 minutes at 4^0^C (acceleration 5 and deacceleration 4). Supernatant was discarded gently and 50µl of staining buffer was added to each well and cells were mixed gently. Surface antibody mixture (anti-CD3-APC 5µl/reaction, anti-CD4-BB515 2µl/reaction, anti-CD8-APC-H7 3µl/reaction, anti-CD25-PE Cy7 2µl/reaction, anti-CD69-PerCP Cy5.5 2.5µl/reaction, anti-CD45Ra-APC-H7 3µl/reaction, anti-iNKT-PE 2µl/reaction) was then added in each well. After gentle mixing, the plate was incubated for 30 minutes at 4^0^C. After incubation cells were washed by adding 100µl of staining buffer in each well and the plate was centrifuged as described earlier. The supernatant was discarded and cells were fixed by adding 100µl 1X FOX P3 buffer in each well and plate was incubated for 30 minutes at 4^0^C. Cells were then washed by adding 100µl of staining buffer in each well. The plate was centrifuged and the supernatant was discarded gently. 150µl of perm wash buffer was then added in each well. After gentle mixing, the plate was incubated at 25^0^C in dark for 30 minutes and centrifuged further for 200g for 3 minutes at 4^0^C. The supernatant was discarded after centrifugation. For intracellular cytokine staining, cells were stained with anti-IL-2-PE Cy7, anti-IFN-γ-PE, anti-IL-4-PE, Cy7, anti-IL-10-PE in each well. The plate was incubated at 4^0^C in dark for 30 minutes. After incubation cells were washed by adding 100µl perm buffer in each well and then centrifuged at 200 g for 3 minutes at 4^0^C. The supernatant was discarded after centrifugation and washing was done by adding 150µl staining buffer in each well and the plate was again centrifuged. The supernatant was discarded and the cells were suspended in staining buffer (200µl/well) for acquisition on flow cytometer. Stained cells were acquired in BD FACS Verse using manufactures protocol and cells were counted using FACS DIVA software. The data were acquired in FACSVerse and analysis was performed on FlowJo version 9.4.4, TreeStar, Inc/FACS Diva software.

### Nitric Oxide Assay

To access the impact of these peptide on activation of monocytic cells, NO titre was quantified using griess regent method. For that purpose, 5 μl of the Nitrate Reductase mixture containing 5 μl of the enzyme cofactor was added to each supernatant. The plates were incubated at room temperature for 1 hr to convert nitrate to nitrite. After that 5μl of the enhancer was added to each well and the plate was incubated further for 10 min. Subsequently 50μl of Griess Reagent R1 and R2 was to each well for 10 min at room temperature for colour development and OD was taken. The pate was read at 540 nm.

## Results

To address the key question of the study, we synthesized peptide using Bioinformatics and homology approach as per scheme mentioned in **(Figure 1).** Based on high homology with MHC –class I peptide, we screened out few peptides mentioned in **Table 1** and selected RV2588c and Rv0148 for our analysis. For the entire study several gating strategies were employed. For quantifying CD3/CD4+, CD45Ra, CD25, CD69 & INKT cells **(Fig S1),** Th-1 response by quantifying IFN/ IL-2/ GMCSF+ CD4/CD8+ T cells (**Fig S2)** and Th-2 response by quantifying IL-4/IL-10+ CD4/CD8+ T cells **(Fig. S3)**. As expected both RV2588c and 0148 activated T cell populations which was revealed the increased number of CD69+ CD4+ (**Fig. 2a**) CD45Ra+ T cell population **(Fig. 2b)** in the PBMC cultures from healthy donors. This increase was much more pronounced in PTB patients indicating recall response in the PBMC from PTB patients. Although Rv0148 enhanced the percentage of iNKT cells also **(Fig 2c)** but it was marginal in the PBMC from healthy donor. However, Rv0148 enhanced these populations 15 times in the PBMC cultures of PTB patients reflecting tuberculosis-associated immune reconstitution inflammatory syndrome (TB-IRIS) in the PBMC of PTB patients. A successful vaccine candidate, apart from enhancing proliferation of effector immune cell population, must trigger an immunogenic response also. To that purpose, Th1 response was evaluated by quantifying **IFN-γ/IL-2+ CD4+ & CD8+ T** cells population **(Fig 3)** in the PBMCs of PTB and Healthy individuals In line with cell proliferation, Rv2588c and Rv0148 enhanced the expression of both **IFN γ** on **CD4+/CD8+T** cells population **(Fig 3A**) in the PBMC from both healthy donors and PTB patients. Interestingly these peptides enhanced the granulocyte-macrophage colony-stimulating factor **(GM-CSF)** producing T cells population **(Fig 3B)**. which are potentially involved in controlling *M. tuberculosis* burden in the PBMC culture of healthy donor. However, these peptides enhanced these populations only marginally in the PBMC from PTB patients indicating their tolerance. A similar response of these peptides was observed on the population of **IL-2** expressing CD4/8+ T cells also **(Fig 3C)** which are important for cell mediated immunity and granuloma formation in PTB patients.

**Figure.2.**
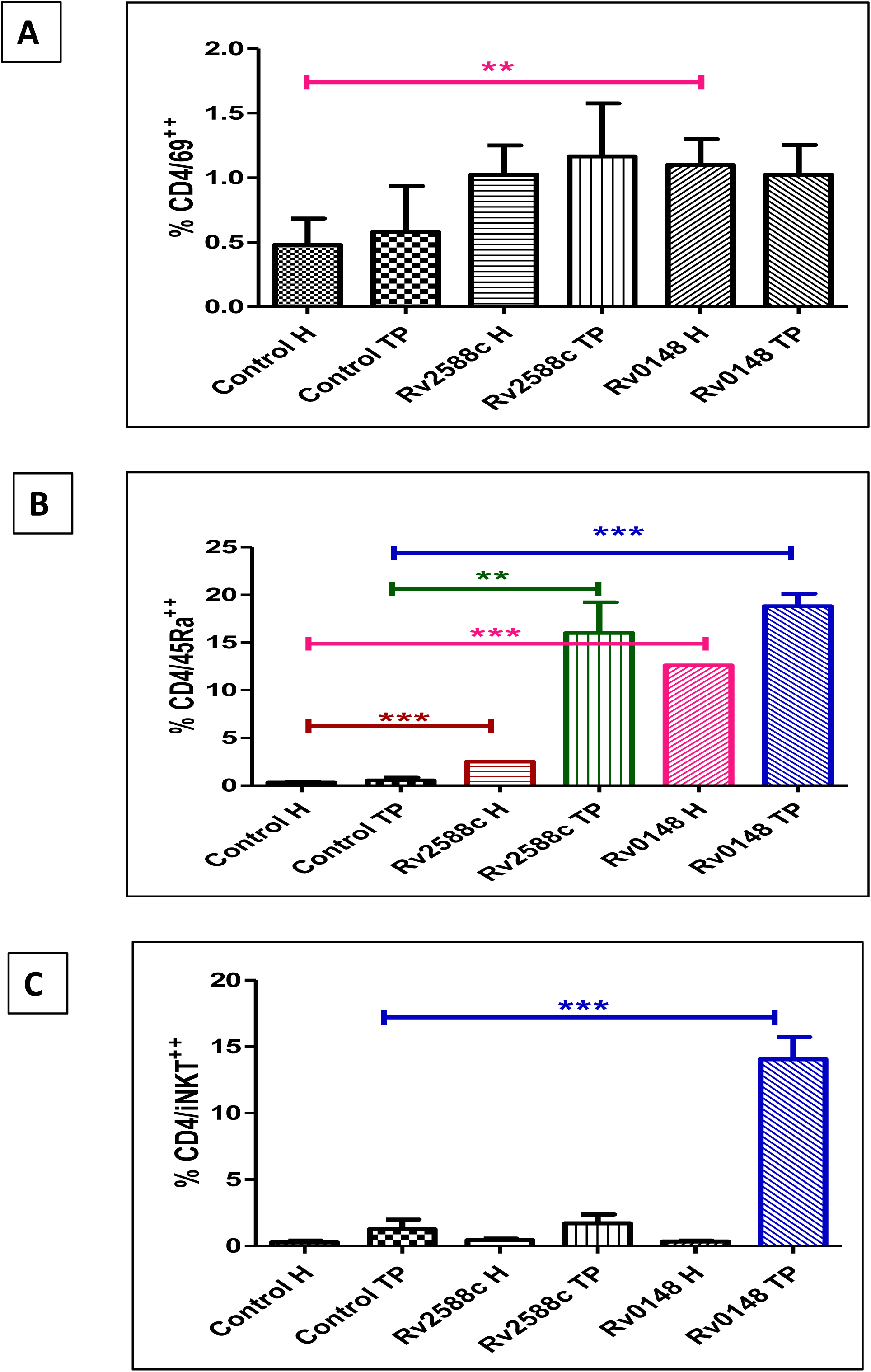
Surface marker expression in response to novel peptide stimulation in TB patients (denoted as TP) and healthy controls (denoted as. **H)** The expression of surface markers on T cells in response to peptide stimulation in peripheral blood samples from TB patients and healthy controls (n=6). Flow cytometry was used to analyse the expression of surface markers (CD25, CD69, CD45Ra, iNKT cells) after stimulation with Rv2588c and Rv0148. Results are expressed as the percentage of positive cells, with data presented as mean ± standard deviation (SD) for each group. Statistical significance was determined using unpaired t-test (Mann-Whitney U test), with p < 0.05 considered significant.

**Figure.3 (A, B, C):**
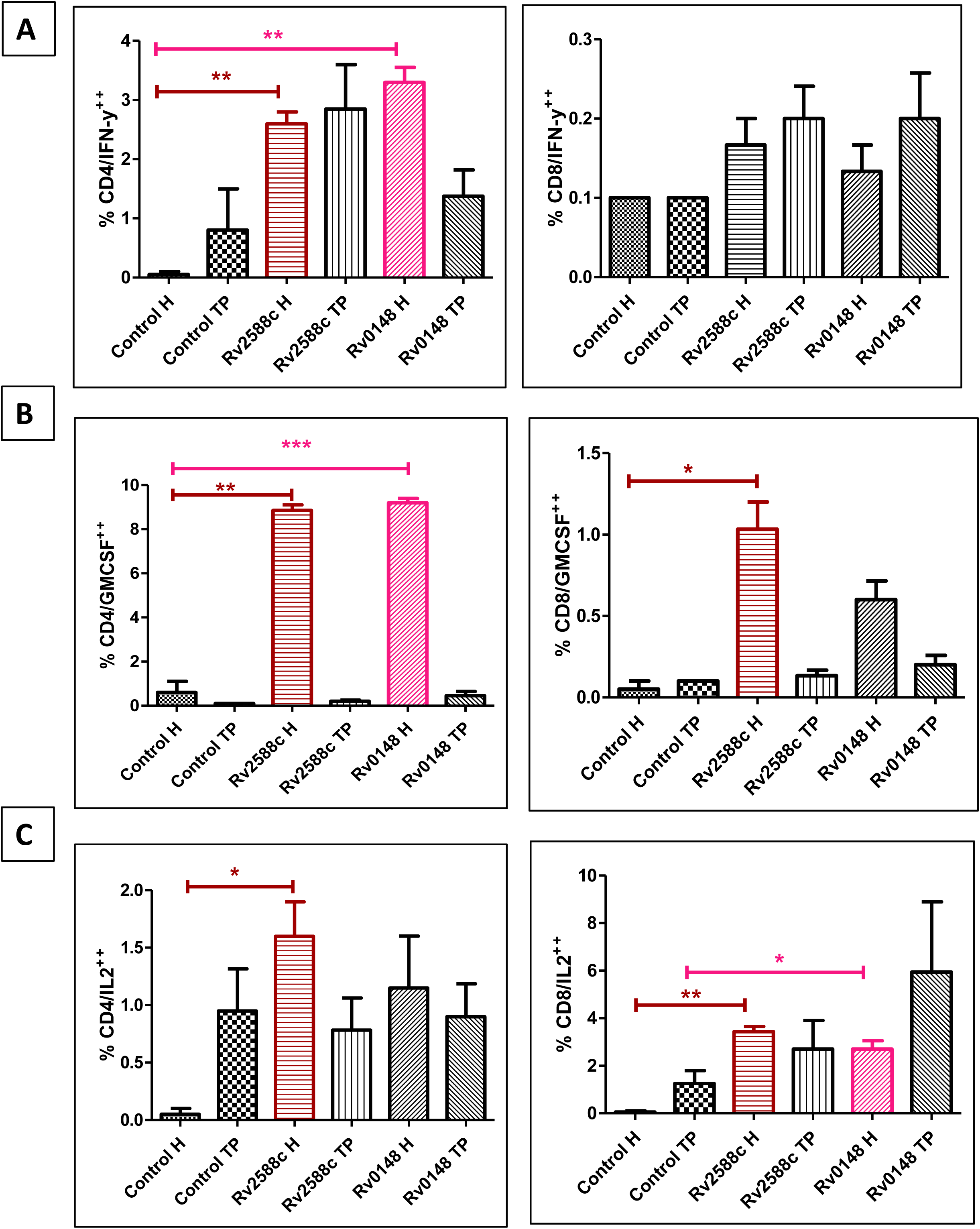
Analysis of Th1 response in TB patients and healthy controls by flow cytometry. The frequency and intensity of Th1 cells (characterized by CD4+ T cells producing IL-2, IFN-γ and GMCSF) in peripheral blood samples from TB patients and healthy controls (n=6). Flow cytometry was used to assess the expression of IFN-γ, a hallmark of Th1 cells, following stimulation with novel peptides Rv2588c and Rv0148. Results are presented as the percentage of IL-2, IFN-γ and GMCSF CD4/8+ T cells. Data are expressed as mean ± standard deviation (SD). Statistical significance was determined using unpaired t-test (Mann-Whitney U test), with p < 0.05 considered significant.

**Table.1:**
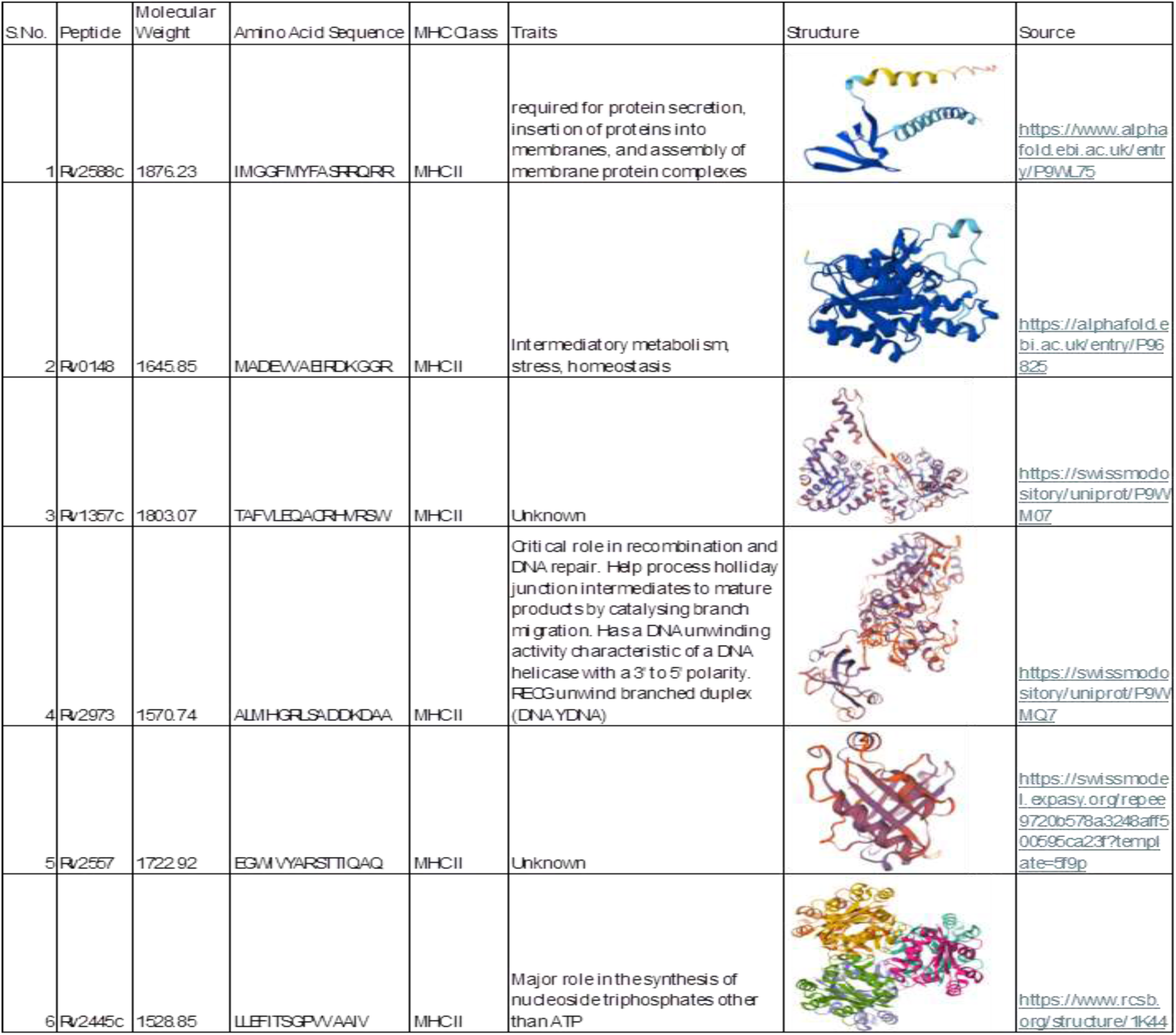
Bioinformatics analysis of MHC-class I restricted peptides from clinical isolate of *M tuberculosis* used in the study. Peptides Rv1357c (TAFVLEQACRHVRSW), Rv2973 (ALMHGRLSADDKDAA), Rv2588C (IMGGFMYFASRRQRR), Rv2557 (EGWIVYARSTTIQAQ), Rv2445c (LLEFITSGPVVAAIV) and Rv2901c (ADAWVWDMYRPARFV) of *M. tuberculosis* having T-cell epitopes as predicted by bioinformatic tools Propred-2, HLApred, HLA-DR4pred for MHC-II and Propred-1, nHLApred, HLApred for MHC-I were synthesized commercially by GL Biochem, Shanghai Ltd. China according to our specification. These peptides have been used in immunoassays to evaluate T cell response in TB patients versus healthy controls.

Any vaccine candidate or adjuvant should not trigger hyperactivation of immune response and should be well tolerable to the recipient. Since Rv2588c and Rv0148 induced protective immune response in the PBMC overlooking Th2 / tolerogenic immune response was formidable for accessing their safety on immune hyperactivation. Therefore we analyzed Suppressor T cells (CD4/25+) T cells population as well as IL-4/ IL-10 expressing CD4+T cell populations which are involved in Th2 response. In line with our presumption both peptides were tolerogenic to the T cell as both these enhanced the number of CD25+CD4+ T cell populations **(Fig 4A).** Apart from IL-4+ CD4+ T cells, the number of Th2 effector T cells population remained significantly less in PTB PBMC indicating ongoing Th1 bias in these cells. Interestingly Rv2588c and Rv0148 enhanced the population of both IL-4 / 10 expressing CD4/8+ T cell populations in healthy PBMC over counter cultures from PTB patients indicting homeostasis in healthy PBMC **(Fig. 4B & C)**.

**Figure.4.**
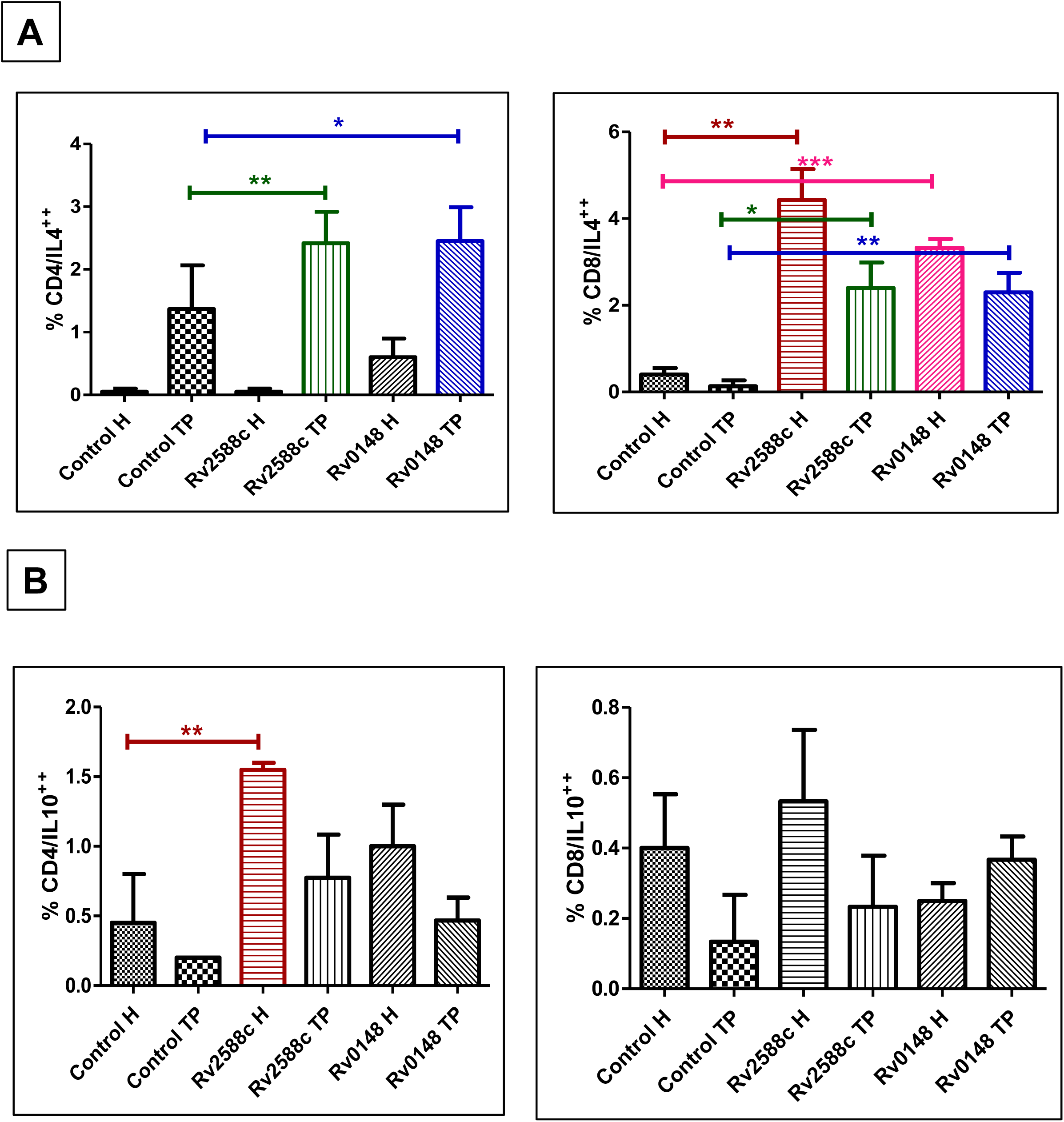
Analysis of Th2 response in TB patients and healthy controls by flow cytometry. The frequency and intensity of Th2 cells (characterized by CD4+ T cells producing IL-4 and IL-10) in peripheral blood samples from TB patients and healthy controls (n=6). Flow cytometry was used to assess the expression of Th2 cytokines following stimulation with novel peptides Rv2588c and Rv0148. Results are presented as the percentage of IL-4/IL-10+ CD4/8+ T cells. Data are expressed as mean ± standard deviation (SD). Statistical significance was determined using unpaired t-test (Mann-Whitney U test), with p < 0.05 considered significant.

Any vaccine candidate and / or immunogen tweaking immunogenic or tolerant immune response should triggers either Th1 or Th2 response. Based on immunomodulatory potential of RV2288c and Rv0148 peptide, we anticipated temporal change in the pattern of Th1/ Th2 cytokines in the culture supernatant of peptide pulsed PBMC. Using sandwich ELISA, we measured several key cytokines which are involved in the immuno-pathological response of TB diseases. Out of Rv2588 and Rv0148, Rv0148 could not modulate the secretion of Th1/2 effector cytokines. Although Rv2588c could not influence the secretion of IFN-γ **(Fig. 5A)** in either of PBMC cultures, it reduced the secretion of IL-4 **(Fig. 5B)** but it remained insignificant. Interestingly Rv2588c enhanced the secretion of IL-1β **(Fig. 5C)** and IL-10 **(Fig 5D)** from the PBMC of PTB patients. These results indicated tolerant nature of Rv2588c peptide in the PBMC.

**Figure.5.**
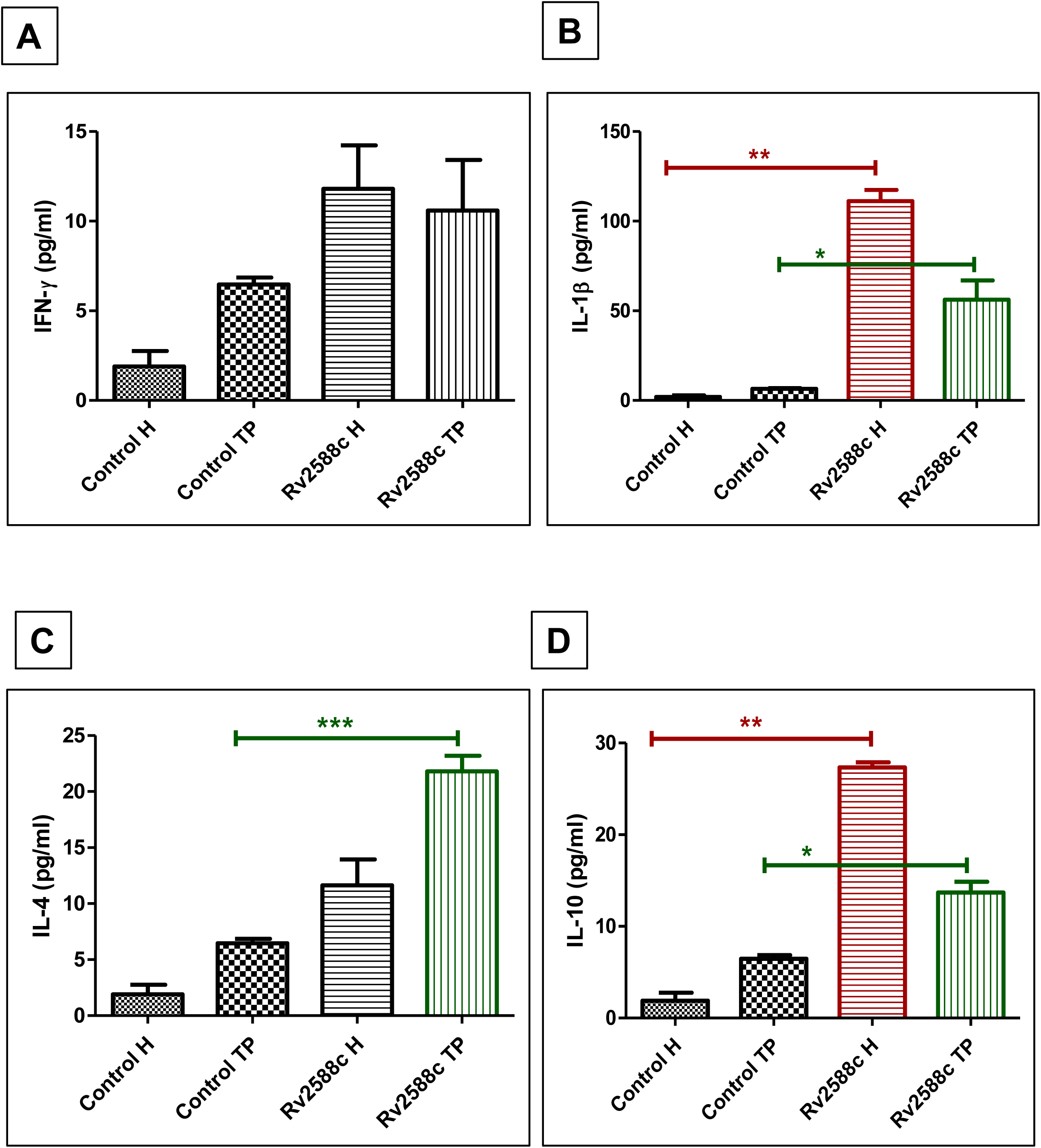
Quantification of Th1 and Th2 levels in culture supernatant from TB patients and healthy controls using sandwich ELISA. The levels of (IFN-γ, IL-1β, IL-4 and IL-10) (measured in pg/ml) against the novel peptides measured by sandwich ELISA in culture supernatant from TB patients and healthy controls (n=10). Results are expressed as mean ± standard deviation (SD) for each group. Statistical significance was determined using one-way ANOVA (non-parametric), with p < 0.05 considered significant.

Based on the Rv2588c/0148 induced T cells priming and Th2 bias, we were keen to analyze whether these T cell restricted epitope would influence monocyte response or not. For that purpose, we analyzed Nitric oxide levels which are mainly secreted by activated monocytes. As expected, these peptides enhanced NO levels in the cultures supernatant of Healthy PBMCs indicating their immune adjuvant nature for monocyte populations **(Fig 6).**

**Fig.6:**
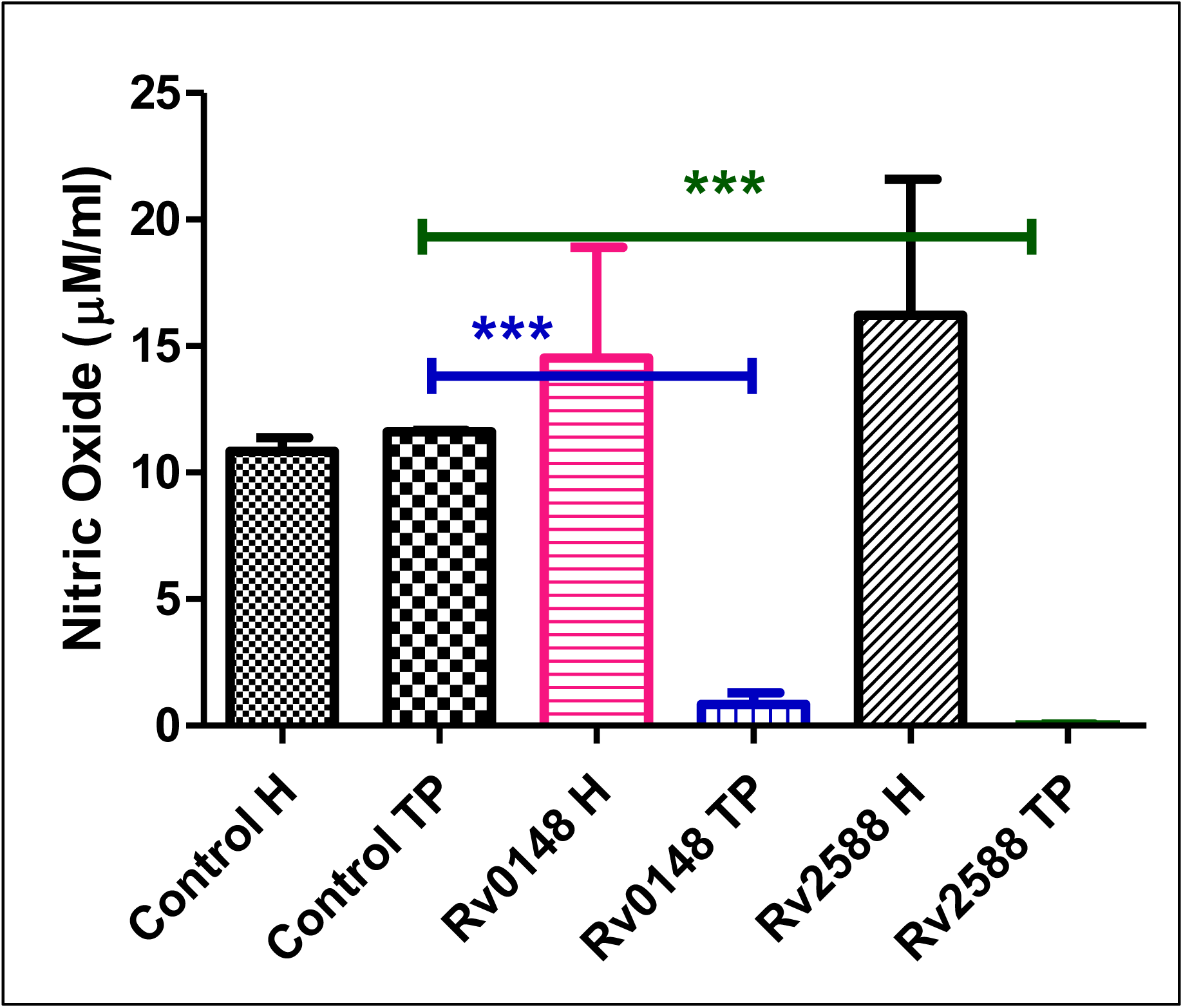
Nitric oxide levels in culture supernatant from TB patients versus healthy controls. The NO titres were quantified and shown here is the mean ± standard deviation (SD) of nitric oxide levels (measured in μM) in supernatant samples from TB patients compared to healthy controls (n=10). Statistical significance was assessed using one-way ANOVA (non-parametric), with p < 0.05 considered significant.

## Discussion

Effective immune response against infectious diseases or tumors depends upon the effective presentation of antigens by APC to T cells which are terminal cells involved in the killing of pathogens or tumors. TB patients harbour heterogeneous lung microenvironments which are characterized by pulmonary fibrosis / caseous granuloma etc. in EPTB or non-reactive TB cases. Such aberrant immuno-pathological micromilieu of lungs adversely impact antigen presentation ability of APC to T cells and impairs their effective activation. Necrosis or *in-situ* polarization / immaturation of APCs are the most common reasons of poor antigen presentation in MDR or refractory cases of TB (which are not supporting treatment regime) displaying high grade pulmonary fibrosis. This is manifested by deregulated ROS, unfolded proteins responses and / or Th2/ 17 biases which affect immune response adversely. Inefficiency of BCG vaccine represents one such cases in TB-endemic regions [17] where antigen processing / presentation by antigen presenting cells (APCs), is regulated negatively by NTM and latent *M.tb* [18] The major reason for the failure of BCG vaccine latent or MDR cases [19] is due to the poor antigen presentation highlighting the significance on antigen presentation for effective immune response against M.tb [18]. In view of refractory immune landscape of TB-endemic populations, exploration or development of peptide vaccine seems to be effective strategies of augmenting effective immune responses against TB in TB-endemic regions [20]. To circumvent this inevitable issues and equal involvement of both CD4 & CD8+T cells in protective immunity against *Mtb* [21], both CD4 & CD8+ T cells epitopes (Rv2588c and Rv0148) having high affinity binding to HLA class I alleles as well as potential to elicit IFN-γ secretion by effector T cells were synthesized. The most striking findings of the study was the impact of synthetic novel MHC class I restricted T-cell epitope (Rv2588c, Rv0148) on cell mediated and innate immune response in PBMC populations and suggests their potential as vaccine candidate. These peptides could play a role in subunit vaccine aimed at generating Th1 response. Evaluation of fusion antigens for developing a TB vaccine needs to be conducted by combining these peptides. Pooled peptide analysis is required for better understanding. Further work is required to achieve the high efficacy and wide applicability of the peptides. In-vivo evaluation is required for validation. The study evaluates the T cell response to these synthetic peptides in pulmonary TB patients and healthy controls, providing insights into the immune mechanisms that could be harnessed for vaccine development. By identifying peptides that elicit a significant immune response, particularly in TB patients, the study suggests that these peptides could serve as potential memory response triggers and new vaccine candidates. The research uses a combination of bioinformatic predictions, ex vivo analysis, and immunological assays to verify the potential of these peptides as immunogenic epitopes, which is a rigorous approach to candidate vaccine evaluation.

The findings of the study could support the development of a peptide-based subunit vaccine, which could be more targeted and potentially more effective than whole protein-based vaccines, as it focuses on specific immunogenic fragments rather than the entire protein. The study lays the groundwork for further research with a larger sample size, which could validate the findings and lead to the development of a new TB vaccine that could have a significant impact on global TB control and elimination efforts. To the best of our knowledge this is the first immune monitoring protocol describing the impact of synthetic novel MHC class I restricted T-cell epitope (Rv2588c, Rv0148) on cell mediated and innate immune response in PBMC populations and suggests their potential as vaccine candidate. Our data prudently suggested that these peptides could be suitable for designing either subunit / peptide vaccine aimed at generating Th1 response against TB. On the basis fo their respective immune adjuvant potential, these \ peptides can be used either combination also for eliciting a robust effector immune responses in TB patients which require further work for achieving high efficacy and wide applicability of the peptides.

## Acknowledgment

We thank all the study participants and the staff of the S.N. Medical Hospital and ICMR-NJIL&OMD. This work was supported by a grant from the ICMR grants to HP and BJ.

## Legend to Supplementary figures

**Figure.S1:**
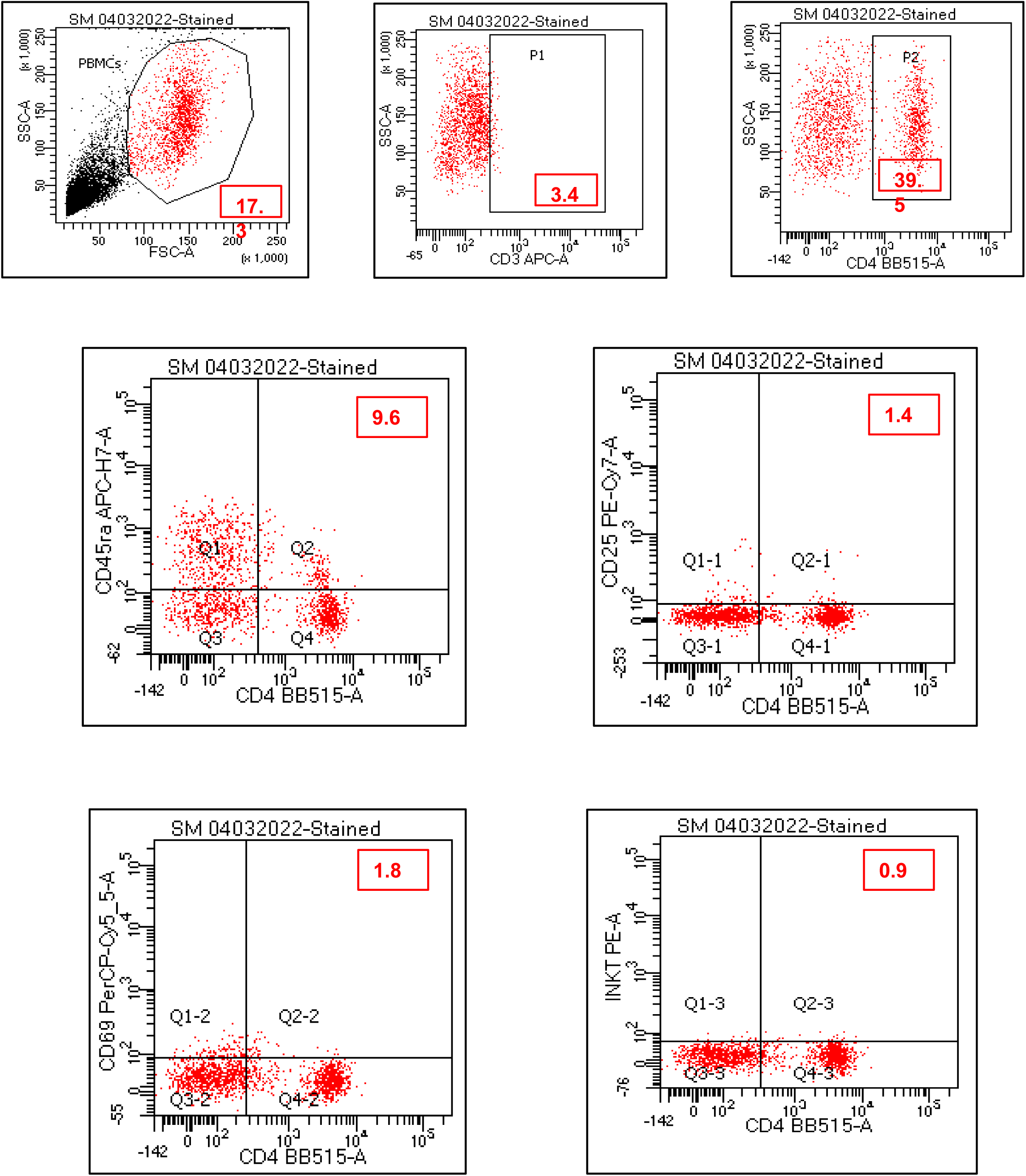
Gating strategy for flow cytometry analysis of (CD3/CD4 positive CD45Ra, CD25, CD69 and iNKT cells) The gating strategy used to identify and analyze (CD3/CD4 positive CD45Ra, CD25, CD69, and iNKT cells). Forward Scatter (FSC) vs. Side Scatter (SSC)-This gate identifies the total population of cells based on size and granularity. Cells are selected within the defined gate to exclude debris and dead cells.

**Figure S2:**
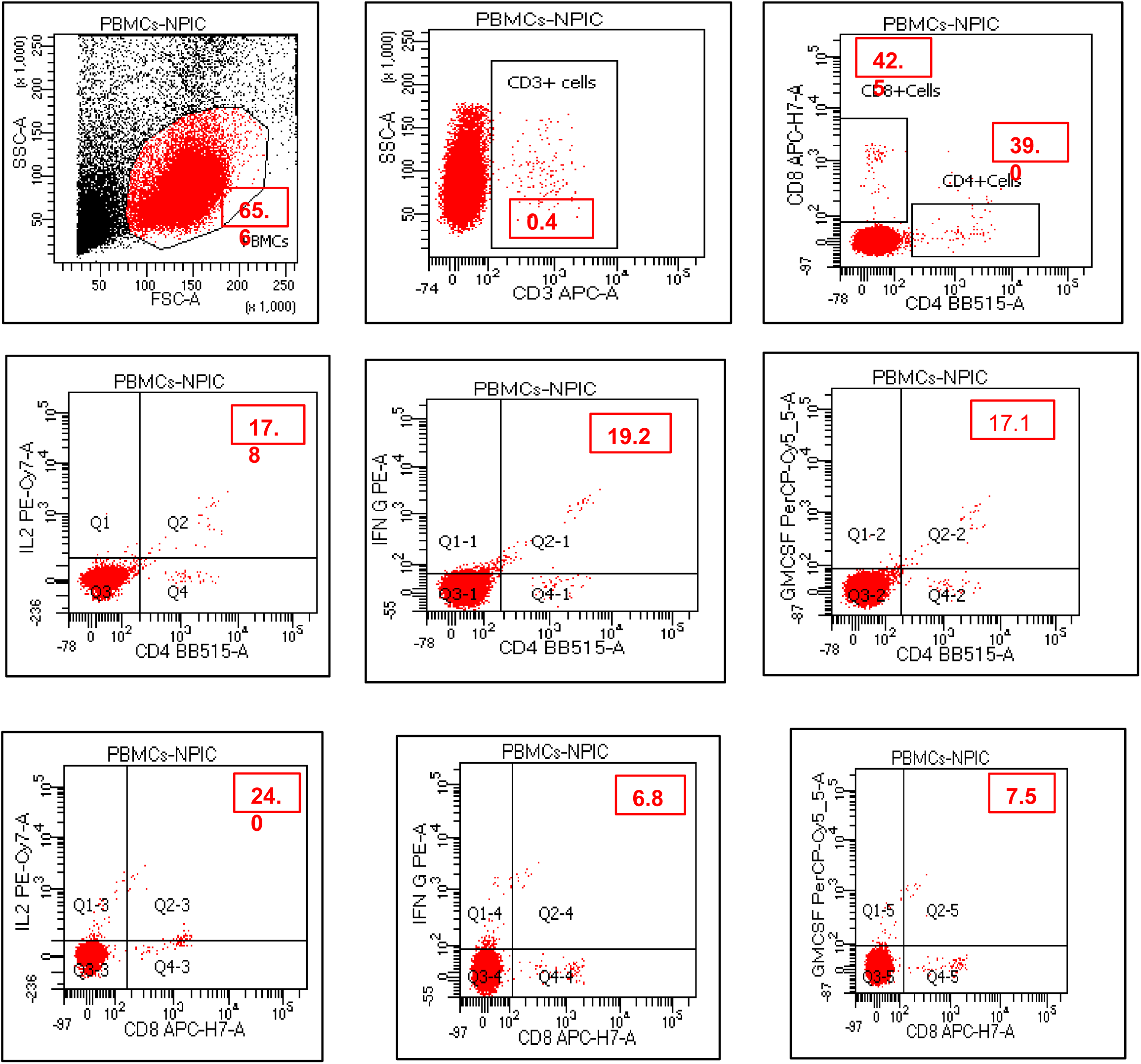
Gating strategy for flow cytometry analysis of Th1 (CD4+/8+ IL-2, IFN-gamma, GMCSF) This figure illustrates the gating strategy used to identify and analyze Th1 (CD4+/8+ IL-2, IFN-gamma, GMCSF). Forward Scatter (FSC) vs. Side Scatter (SSC)-This gate identifies the total population of cells based on size and granularity. Cells are selected within the defined gate to exclude debris and dead cells.

**Figure S3:**
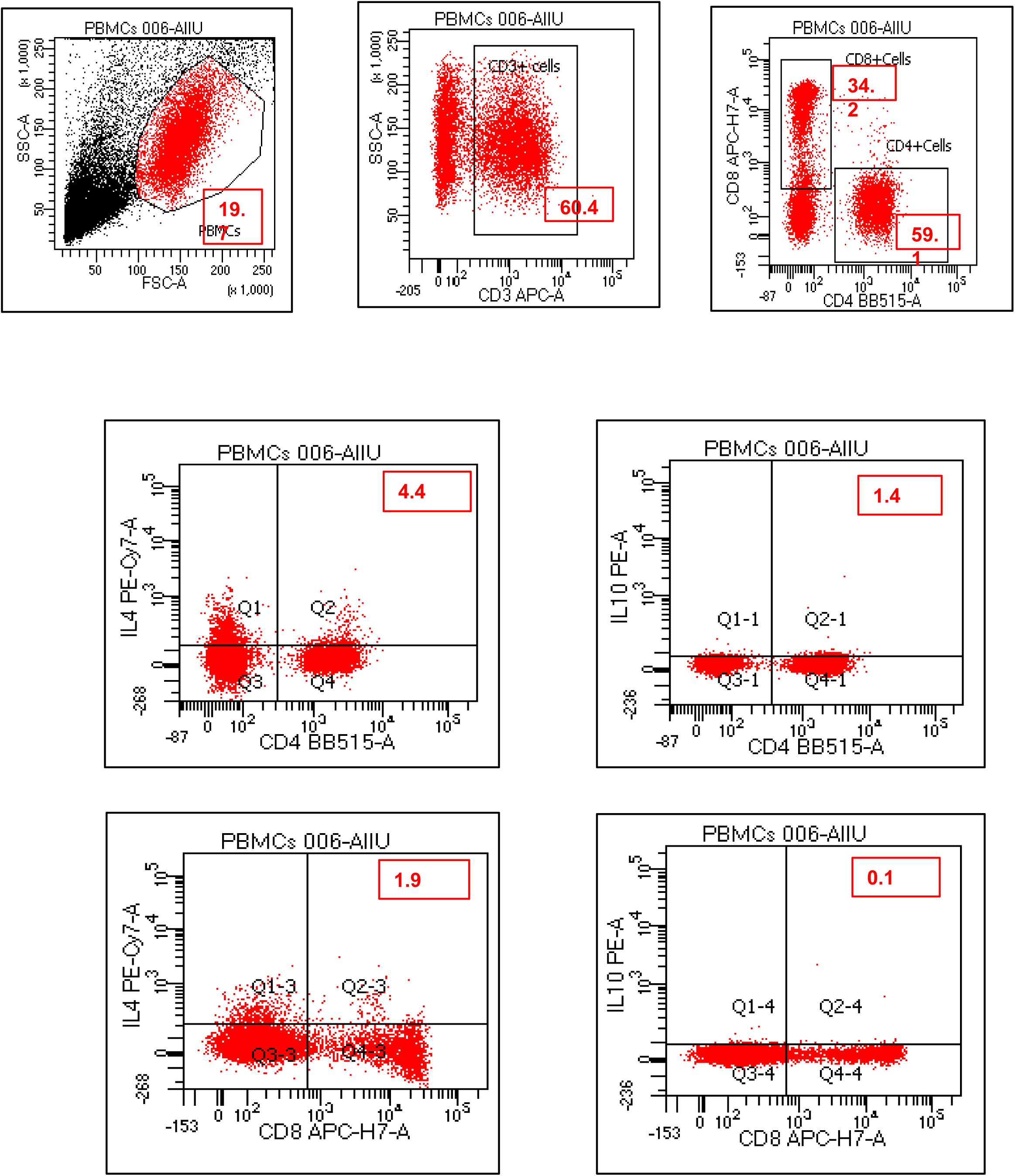
Gating strategy for flow cytometry analysis of Th2 (CD4+/8+ IL-2, IL-10) This figure illustrates the gating strategy used to identify and analyze Th2 (CD4+/8+ IL-2, IL-10). Forward Scatter (FSC) vs. Side Scatter (SSC)-This gate identifies the total population of cells based on size and granularity. Cells are selected within the defined gate to exclude debris and dead cells.

**Figure. S4:**
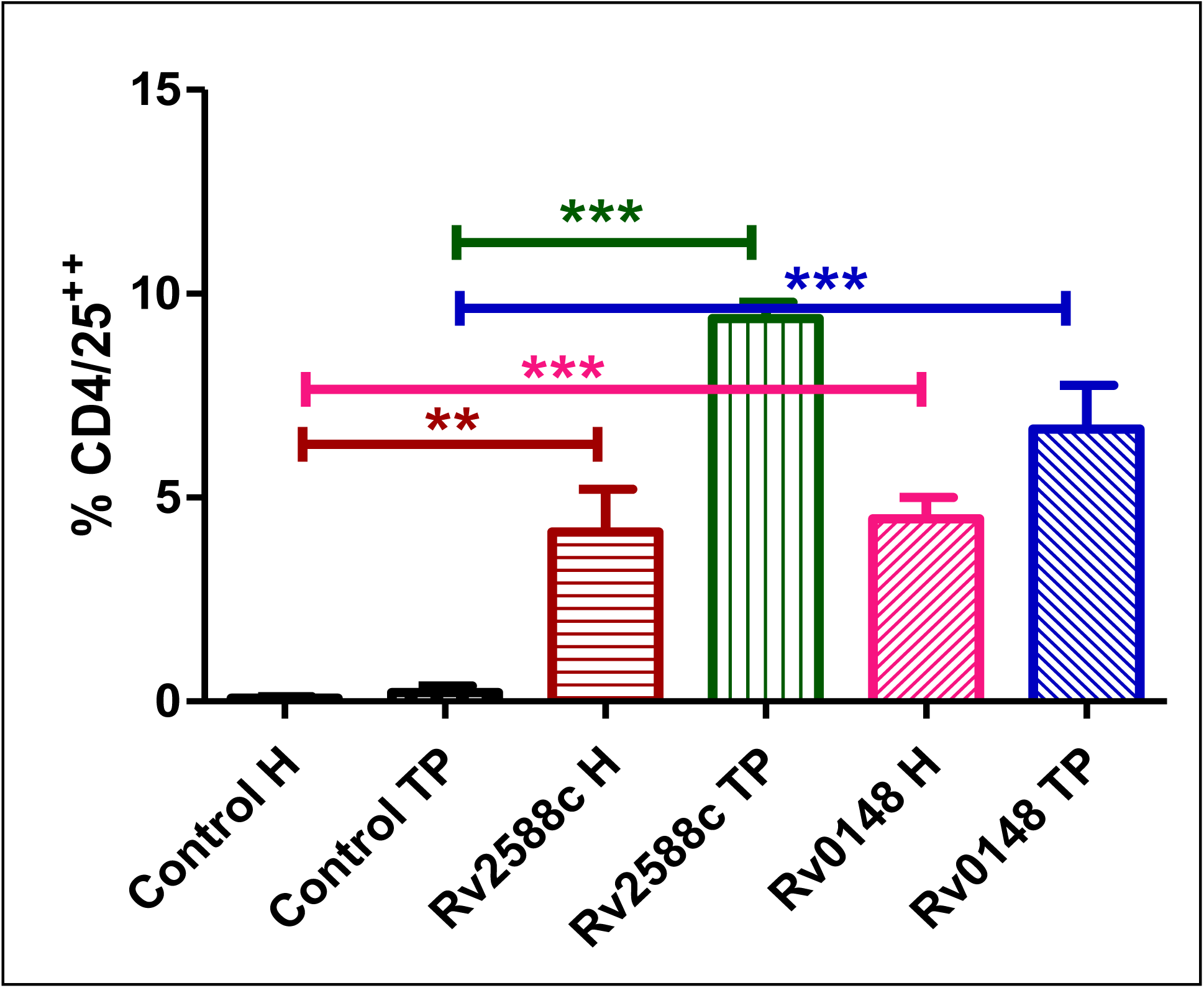
Assessment of suppressor T cells in the PBMC. Flow cytometry was used to analyse the expression of surface markers (CD25) after stimulation with Rv2588c and Rv0148. Results are expressed as the percentage of positive cells, with data presented as mean ± standard deviation (SD) for each group. Statistical significance was determined using unpaired t-test (Mann-Whitney U test), with p < 0.05 considered significant.

## References

1. Goletti, D., et al., World Tuberculosis Day 2023 theme “Yes! We Can End TB!”. International Journal of Infectious Diseases, 2023. 130: p. S1–S3.

2. Geluk, A., et al., Innovative strategies to identify M. tuberculosis antigens and epitopes using genome-wide analyses. Frontiers in immunology, 2014. 5: p. 256.

3. Kaufmann, S.H., J. Weiner, and C.F. von Reyn, Novel approaches to tuberculosis vaccine development. International Journal of Infectious Diseases, 2017. 56: p. 263–267.

4. Li, W., et al., Peptide vaccine: progress and challenges. Vaccines, 2014. 2(3): p. 515–536.

5. Skwarczynski, M. and I. Toth, Peptide-based synthetic vaccines. Chemical science, 2016. 7(2): p. 842–854.

6. Kornfeld, H., G. Mancino, and V. Colizzi, The role of macrophage cell death in tuberculosis. Cell Death & Differentiation, 1999. 6(1): p. 71–78.

7. Stenger, S., Cytolytic T cells in the immune response to mycobacterium tuberculosis. Scandinavian journal of infectious diseases, 2001. 33(7): p. 483–487.

8. Gideon, H.P. and J.L. Flynn, Latent tuberculosis: what the host “sees”? Immunologic research, 2011. 50: p. 202–212.

9. Kaufmann, S.H., The macrophage in tuberculosis: sinner or saint? The T cell decides. Pathobiology, 1991. 59(3): p. 153–155.

10. Philips, J.A. and J.D. Ernst, Tuberculosis pathogenesis and immunity. Annual Review of Pathology: Mechanisms of Disease, 2012. 7(1): p. 353–384.

11. Mogues, T., et al., The relative importance of T cell subsets in immunity and immunopathology of airborne Mycobacterium tuberculosis infection in mice. The Journal of experimental medicine, 2001. 193(3): p. 271–280.

12. Cooper, A.M., et al., Disseminated tuberculosis in interferon gamma gene-disrupted mice. The Journal of experimental medicine, 1993. 178(6): p. 2243–2247.

13. Sud, D., et al., Contribution of CD8+ T cells to control of Mycobacterium tuberculosis infection. The Journal of Immunology, 2006. 176(7): p. 4296–4314.

14. Kumar, G., et al., Proteomics of culture filtrate of prevalent Mycobacterium tuberculosis strains: 2D-PAGE map and MALDI-TOF/MS analysis. SLAS DISCOVERY: Advancing Life Sciences R&D, 2017. 22(9): p. 1142–1149.

15. Kumar, G., et al., *Whole cell &* culture filtrate proteins from prevalent genotypes of Mycobacterium tuberculosis provoke better antibody & T cell response than laboratory strain H37Rv. Indian Journal of Medical Research, 2012. 135(5): p. 745–755.

16. Sharma, N., et al., Immunological depiction of synthetic B-cell epitopes of Mycobacterium tuberculosis. The International Journal of Mycobacteriology, 2023. 12(4): p. 380–387.

17. Moliva, J.I., J. Turner, and J.B. Torrelles, Immune responses to bacillus Calmette–Guérin vaccination: why do they fail to protect against Mycobacterium tuberculosis? Frontiers in immunology, 2017. 8: p. 407.

18. Gowthaman, U., et al., Lipidated promiscuous peptides vaccine for tuberculosis-endemic regions. Trends in Molecular Medicine, 2012. 18(10): p. 607–614.

19. Dockrell, H.M. and S.G. Smith, What have we learnt about BCG vaccination in the last 20 years? Frontiers in immunology, 2017. 8: p. 1134.

20. Gowthaman, U., et al., Promiscuous peptide of 16 kDa antigen linked to Pam2Cys protects against Mycobacterium tuberculosis by evoking enduring memory T-cell response. Journal of Infectious Diseases, 2011. 204(9): p. 1328–1338.

21. Ritz, N., et al., Comparable CD4 and CD8 T cell responses and cytokine release after at-birth and delayed BCG immunisation in infants born in Australia. Vaccine, 2016. 34(35): p. 4132–4139.

